# Motor imagery training is beneficial for motor memory of upper and lower limb tasks in very old adults

**DOI:** 10.1101/2022.10.04.510767

**Authors:** Pauline M Hilt, Mathilde Bertrand, Léonard Féasson, Florent Lebon, France Mourey, Célia Ruffino, Vianney Rozand

## Abstract

Human aging is associated with a decline in the capacity to memorize recently acquired motor skills. Motor imagery training is a beneficial method to compensate for this deterioration in old adults. It is not yet known whether these beneficial effects are maintained in very old adults (>80 years), more affected by the degeneration processes. The aim of this study was to evaluate the effectiveness of a mental training session of motor imagery on the memorization of new motor skills acquired through physical practice in very old adults. Thirty very old adults performed 3 actual trials of a manual dexterity task (session 1) or a sequential footstep task (session 2) as fast as they could before and after a 20-min motor imagery training (mental-training group) or watching a documentary for 20 min (control group). Performance was improved after 3 actual trials for both tasks and both groups. For the control group, performance decreased in the manual dexterity task after the 20-min break and remained stable in the sequential stepping task. For the mental-training group, performance was maintained in the manual dexterity task after the 20-min motor imagery training and increased in the sequential stepping task. These results extended the benefits of motor imagery training to the very old population, showing that even a short motor imagery training improved their performance and favor the motor memory process. These results confirmed that motor imagery training is an effective method to complement traditional rehabilitation protocols.

## INTRODUCTION

Aging is associated with progressive neural degenerative processes inducing impairments in cognitive function (Andrews-Hanna et al., 2007) such as declarative (LaVoie and Cobia, 2007) or motor (Robertson et al., 2004) memory. Motor memory, or memorization, is involved in motor learning through the encoding of planning details of the new-learned movement (Classen et al., 1998). Although conflicting results exist in the literature on the age-related impairment in motor learning capacities (Anguera et al., 2011; Huang and Ahmed, 2014), motor memory of new-learned movements is altered in the elderly after a 30-min break on a dexterity manual task (Ruffino et al., 2019) and after a 5-min break on a walking perturbation task (Malone and Bastian, 2016). Specifically, old adults showed a deterioration of performance after the resting period whereas young adults preserved their performance compared to the performance measured before the break (Malone and Bastian, 2016; Ruffino et al., 2019).

Although pharmacological interventions can reduce the age-related deterioration in motor memory (Flöel et al., 2005), non-medical methods such as motor imagery may be a relevant alternative to limit the use of drugs. Motor imagery is the mental simulation of a movement without overt execution (Jeannerod, 1994). Mental representation of an action shares similar neural network with those activated during physical practice (Decety et al., 1994; Gerardin et al., 2000), like motor-related regions (Guillot and Collet, 2008; Guillot et al., 2009). A large body of literature showed that motor imagery training over one or repeated sessions was effective to improve motor performance such as maximal strength (Yue and Cole, 1992; Zijdewind et al., 2003) or motor skills (Feltz and Landers, 1983; Gentili et al., 2006). In old adults, the ability to imagine simple movements is preserved but become more difficult for complex or unusual movements (Saimpont et al., 2013). Hence, motor imagery training is still effective to improve motor performance in old adults (Hamel and Lajoie, 2005; Ruffino et al., 2019). Indeed, (Ruffino et al., 2019) showed that old adults who performed a motor imagery training for 30 min preserved the gains obtained on a dexterity manual task, whereas those who rested for 30 min erased these gains. Whether these beneficial effects of motor imagery on motor memory are also maintained in very old adults (>80 years) remains unknown. Yet, very old adults are largely affected by the deterioration process of neuromuscular function (Varesco et al., 2022) and would greatly benefit from this kind of rehabilitation technic. Although the temporal congruence between actual and imagined movements remains similar in young and old adults, imagined movement duration seems altered at very old age (>80 years; Schott and Munzert, 2007). Similar results have been observed for old (62-67 years) and older individuals (71-75 years), with only the latter group showing significant differences in performance compared to young adults (Skoura et al., 2005). These results suggest that the beneficial effects of motor imagery on motor skill memorization may be lowered in very old adults.

Thus, the aim of the present study was to evaluate the effectiveness of a single session of mental training, using motor imagery, on memorization of motor skill acquired by physical practice in very old adults. As in the study of Ruffino et al. (2019), we tested this memorization process with a dexterity manual task (upper limbs). Due to the rapid deterioration of functional capacities in very old adults, notably in balance and mobility functions (Studenski et al., 2011), we also tested the effects of mental training on a sequential footstep task (lower limbs). We hypothesized that motor imagery would have a positive effect on motor memory for both upper and lower limb tasks, but lower than that previously observed in old adults (Ruffino et al., 2019).

## METHODS

### Participants

Thirty right-handed healthy very old adults (19 women, age: 86 ± 2 years, height: 163 ± 8 cm, weight: 62 ± 12 kg, Mini-Mental Stage Examination (Folstein et al., 1975) = 29 ± 1, range [22 - 30]) volunteered to participate in the experiment and were clearly informed on the experimental procedures prior giving their written consent. Participants had no neurological or physical disorders or disabilities. All the experimental procedures were approved by the French Ethics Committee Sud-Ouest et Outre-Mer 1 and were performed in accordance with the declaration of Helsinki, except for registration in a database (ClinicalTrials.gov, Identifier: NCT04018196).

### Experimental procedures

Participants, randomly assigned to a mental-training (n = 15) or a control (n = 15) group, visited the laboratory on two occasions separated by 24 hours. During the first visit, participants completed an imagery ability questionnaire (mental-training group only) and functional tests (both groups, see below for details). Then, we assessed the effects of motor imagery training on a dexterity manual task. The participants were comfortably seated on a chair placed at 20 cm in front of a table to perform a modified version of Nine Hole Peg Test (NHPT; Figure 1). The original NHPT has been modified to increase the difficulty, duration and number of movements (Ruffino et al., 2019). The NHPT required the participants moving the 9 sticks as fast as possible into 9 holes in a pre-determined order and then replacing them in a box. They started moving the stick from hole 1 to hole A, then from hole 2 to hole B, and so on. Once all sticks were placed into the corresponding holes, the sticks were moved into the box in the same order. The experimenter started the timer when the participant touched the first stick and stopped it when the last stick was in the box. Both groups performed 3 trials (a total of 108 movements) with 1-min rest between each trial at PreTest and PostTest. After performing the 3 PreTest trials, the participants either watched a non-emotional documentary for 20 min (“Home”, directed by Y. Arthus-Bertrand, 2009; control group) or performed a mental training using motor imagery (mental-training group). The duration of the documentary was similar to the duration of the mental training. The participants of the mental-training group were instructed to imagine themselves performing the NHPT as fast as possible, combining the kinesthetic and visual (first-person perspective) modalities. They performed 3 blocks of 10 trials with 5-s rest between trials and 1-min rest between blocks to avoid mental fatigue (Rozand et al., 2016). After pilot studies, the number of blocks was adjusted compared to a similar study of our research group on young and old adults (Ruffino et al., 2019) because the very old adults had difficulty staying focused during 5 blocks. Therefore, we reduced the number of blocks to 3 during the mental training. The duration of each imagined trial was measured to ensure isochrony between imagined and actual trials. The experimenter started the timer when the participant touched the first stick and stopped the timer when the participant dropped this same first stick.

**Figure 1:**
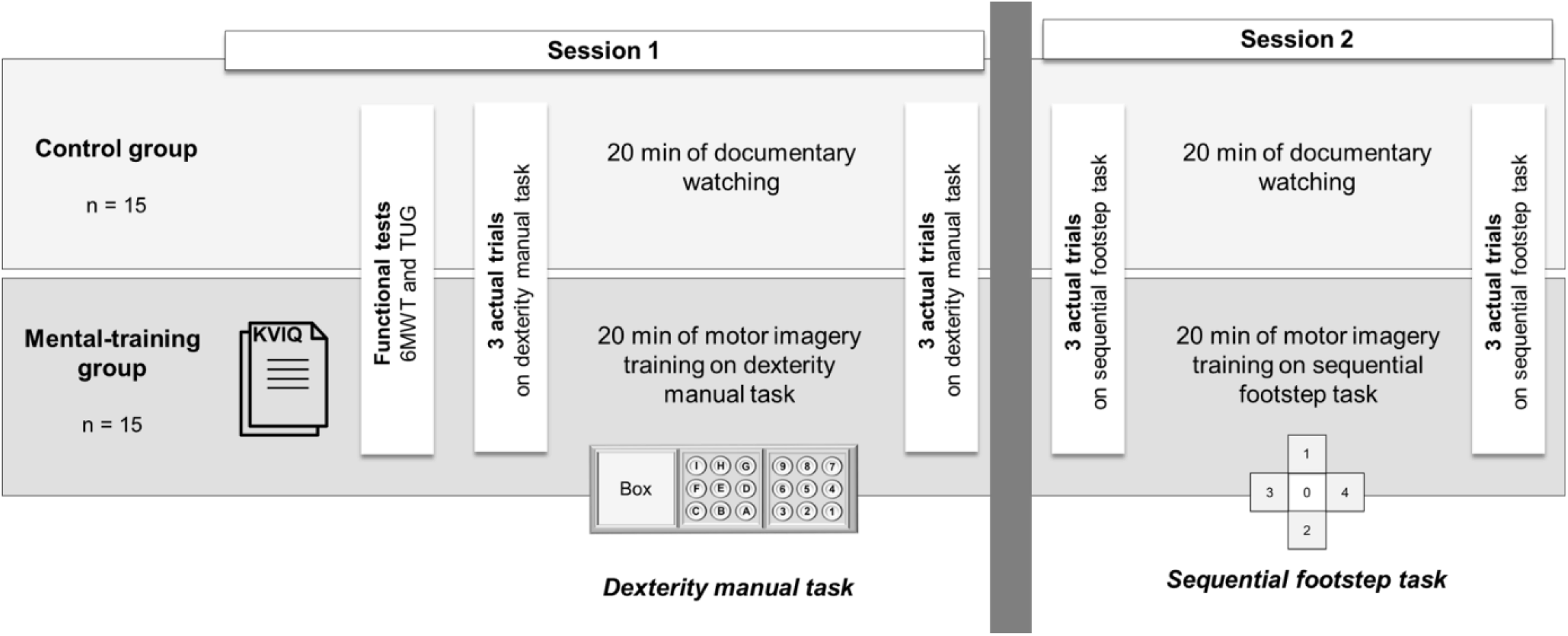
Illustration of the experimental protocol.

During the second visit performed at least 24 h after the first visit, we assessed the effects of mental training on skill performance improvement of lower limbs, using a sequential footstep task. To complete a sequence, the participants started in box 0, moved to box 1, moved back to box 0, then box 2, up to box 4 and moved back to box 0. They were required to move their two feet in each box (surface of 0,25 m^2^), and performed two sequences per trial. The experimenter started the timer when the participant left the first foot and stopped it when the second foot touched the last box at the end of the trial. The participants performed 3 trials as fast as possible with 1-min rest between each trial at PreTest and PostTest. After performing the PreTest, the participants either watched the non-emotional documentary for 20 min (“Home”, directed by Y. Arthus-Bertrand, 2009; control group) or performed a mental training using motor imagery (mental-training group). The participants of the mental-training group were instructed to imagine themselves performing the footstep task as fast as possible, combining the kinesthetic and visual (first-person perspective) modalities. They were sitting on a chair on box 0. They performed 3 blocks of 10 trials with 5-s rest between trials and 1-min rest between blocks (Figure 1). The duration of the imagined trials was assessed to ensure isochrony between imagined and actual trials. The experimenter started the timer when the participant lift their right foot and stopped the timer when the participant put his foot down.

### Imagery ability

Participants of the mental-training group completed the revised version of the Kinesthetic and Visual Imagery Questionnaire (KVIQ) (Malouin et al., 2007) at the beginning of the first visit. The KVIQ consisted of 10 simple movements involving either the upper-limb, lower-limb or the trunk actually performed and then imagined with visual and kinesthetic modalities. After each imagined movement from a first-person perspective, the participants rated on a five-point scale the clarity of images (visual mode) or the intensity of the sensations (kinesthetic mode) from 1 (no image/no sensation) to 5 (image as clear as when seeing the movement/sensation as intense as when executing the movement).

During mental training, the participants rated the vividness of their imagined movements on the same 5-point scale after each block.

### Functional tests

All participants performed the 6-minute Walk Test (6MWT), which consisted of covering the longest distance possible in 6 minutes in a corridor of 30 meters (Brooks et al., 2003). They also performed the Timed-Up-and-Go test (TUG), which required the participants to stand up from a chair, walk a distance of three meters, turn around, return, and sit down on the chair (Mathias et al., 1986). The experimenter started the stopwatch when the back of the participant moved from the chair and stopped it when the participant sat with his/her back on the back of the chair.

### Experiment on old adults

To compare the effect of mental training across ages, we used a dataset already published (Ruffino et al. 2019) using the NHPT task on a group of old adults (n = 13; mean age: 72 ± 4 year-old, 7 females, MMSE = 29 ± 1). Ruffino and collaborators used the same task (modified version of the Nine Hole Peg Test) in the same conditions. Their mental training was however longer (5 blocks of 10 trials). They also included a control group (n = 10; mean age: 74 ± 6 year-old, 9 females, MMSE = 29 ± 1) who watched the same non-emotional documentary (“Home”, directed by Y. Arthus-Bertrand, 2009) for 30 min. To note that old and very old participants had similar performances on the MMSE test (p=0.79, t=0.69).

### Statistical analysis

Shapiro-Wilk test revealed that our dataset was not normally distributed (P<0.05). For this reason, we used multiple two-tail permutation tests (5000 permutations; MATLAB function mult_comp_perm_t1) for all comparisons. All analyses were performed using a custom software written in MATLAB (Mathworks). P-values were corrected for multiple comparisons using the Benjamini-Hochberg false discovery rate (MATLAB function fdr_bh).

Scores of MIQ-R and MMSE, TUG duration, 6MWT distance were compared between the mental-training and control groups, and movement durations were compared between imagined and actual trials in the mental-training group.

Initial performance between the two groups (mental-training vs. control) was compared within each PreTest trial and between the means of the 3 PreTest trials for NHPT and footstep task. Trial-by-trial evolution in the PreTest was evaluated in each group for both tasks.

To test the impact of mental training or the break period on performance retention, we compared the averaged movement duration between PreTest and PostTest, and also between the last PreTest trial (PreTest 3) and the first PostTest trial (PostTest 1) for the NHPT and footstep task. Finally, to evaluate the slope of learning within PreTest and PostTest, we computed linear regressions over the duration of the 3 trials. The slope values were then compared across groups to evaluate potential changes in learning rate after intervention.

Data are presented as mean ± standard deviation in the text and table.

## RESULTS

### Cognitive and functional capacities

Participants of the mental-training and control groups showed no significant difference in MMSE score (p=0.37, t=1.54), TUG duration (p=0.84, t=0.74) and 6MWT distance (p=0.84, t=-0.82; Table 1).

**Table 1.** Cognitive and functional capacities of the mental-training and control groups. MMSE: Mini-Mental State Examination. KVIQ: Kinesthetic and visual imagery questionnaire. 6MWT: 6-min Walk Test. TUG: Timed-Up and Go test.

|  | Mental-training group | Control group |
| --- | --- | --- |
| MMSE | 29 ± 1 | 29 ± 2 |
| KVIQ | 86 ± 15 |  |
| 6MWT (m) | 440 ± 68 | 426 ± 69 |
| TUG (s) | 9.0 ± 1.5 | 9.4 ± 2.0 |

Duration of the imagined movement (average across the 3 blocs; NHPT: 18.7 ± 9.4 s, footstep task: 16.7 ± 7.3 s) was not significantly different than the duration of the actual trials (average across PreTest and PostTest trials; NHPT: 18.2 ± 4.2 s, footstep task: 19.3 ± 5.3 s). Furthermore, the vividness of the imagined movements was close to maximum score (4.67 ±0.42), suggesting a good capacity to imagine.

### Motor performance at PreTests

For the modified version of the NHPT, permutation tests confirmed the absence of significant difference in initial performance between the two groups (PreTest 1: p=0.99, t=-0.11; PreTest2: p=0.99, t=-0.17; PreTest3: p=0.93, t=-0.37; mean PreTest: p=0.84, t=-0.21). Within each group, the analyses revealed a longer duration in PreTest 1 (Mental-training: 20.6 ± 3.9 s; Control: 20.8 ± 4.3 s) compared to PreTest 2 (Mental-training: 19.0 ± 4.1 s, p<0.01, t=-3.65; Control: 19.3 ± 3.5 s, p<0.05, t=-2.54) and PreTest 3 (Mental-training: 17.8 ± 4.1 s, p<0.01, t=-4.10; Control: 18.3 ± 2.5 s, p<0.01, t=-3.44). PreTest 2 was significantly longer than PreTest 3 for the mental-training group only (p<0.01, t=-4.10). The learning’s slopes computed over the 3 PreTest trials were similar between the two groups (Mental-training slope: −1.4 ± 1.0; Control slope: −1.2 ± 1.4; p=0.99, t=-0.24).

For the footstep task, no significant difference was observed in initial performance between the two groups (PreTest 1: p=0.32, t=1.37; PreTest2: p=0.29, t=1.42; PreTest3: p=0.62, t=0.86; mean PreTest: p=0.22, t=1.31). Movement duration was longer in PreTest 1 (Mental-training: 22.7 ± 5.3 s; Control: 20.8 ± 2.9 s) compared to PreTest 2 (Mental-training: 20.6 ± 5.4 s, p<0.05, t=-3.10; Control: 18.9 ± 2.8 s, p<0.001, t=-4.49) and PreTest 3 (Mental-training: 19.8 ± 5.5 s, p<0.01, t=-3.83; Control: 18.6 ± 3.3 s, p<0.05, t=-2.81). No significant difference was found between PreTest 2 and PreTest 3 (Mental-training: p=0.08, t=-2.25; Control: p=0.88, t=-0.50). The learning’s slopes computed over the 3 PreTest trials were similar between the two groups (Mental-training slope: −1.4 ± 1.5; Control slope: −1.2 ± 1.4; p=0.97, t=-0.41).

These results indicated that the two groups improved their performance through the PreTest trials in both upper and lower limb tasks.

### Motor performance following the intervention

Following the PreTest trials, the control group watched a documentary for 20 min to control for the evolution of the memorization process, while the experimental group performed a mental training consisting of 3 blocks of 10 trials (~20 min) to evaluate the effects of motor imagery-based training on the memorization of a new-learned skill.

For the NHPT, the averaged movement duration across the three trials significantly decreased for the mental-training group (PreTest: 19.1 ± 3.9 s, PostTest: 17.2 ± 4.0 s, p<0.001, t=-9.58), while no difference was observed for the control group (PreTest: 19.4 ± 3.2 s, PostTest: 18.5 ± 2.4 s, p=0.30, t=-1.72). When performing a trial-by-trial analysis, movement duration was not significantly different from PreTest3 to PostTest 1 (p=0.88, t=0.72) for the mental-training group, but increased from PreTest3 to PostTest 1 (p<0.05, t=3.06) for the control group (Figure 1). The learning’s slopes computed over the 3 PostTest trials was lower for the mental-training group (−0.8 ± 0.5) than the control group (−1.2 ± 0.9, p<0.05, t=3.26).

These results indicated that motor-imagery training help to prevent the motor memory deficit observed after a short break period for the control group.

For the footstep task, the averaged movement duration significantly decreased from PreTest to PostTest for the mental-training group (PreTest: 21.0 ± 5.2 s, PostTest: 17.5 ± 4.6 s, p<0.001, t=-8.03), while no difference was observed for the control group (PreTest: 19.4 ± 2.7 s, PostTest: 17.9 ± 3.1 s, p=0.07, t=-2.66). When performing a trial-by-trial analysis, movement duration significantly decreased from PreTest 3 to PostTest 1 (p<0.01, t=-4.26) for the mental-training group, but was not significantly different from PreTest3 to PostTest 1 (p=0.79, t=0.91) for the control group (Figure 2). Linear regression over the duration of the 3 PostTest trials was lower for the mental-training group (−0.4 ± 0.6) than the control group (−1.2 ± 0.9, p<0.05, t=3.26).

**Figure 2.**
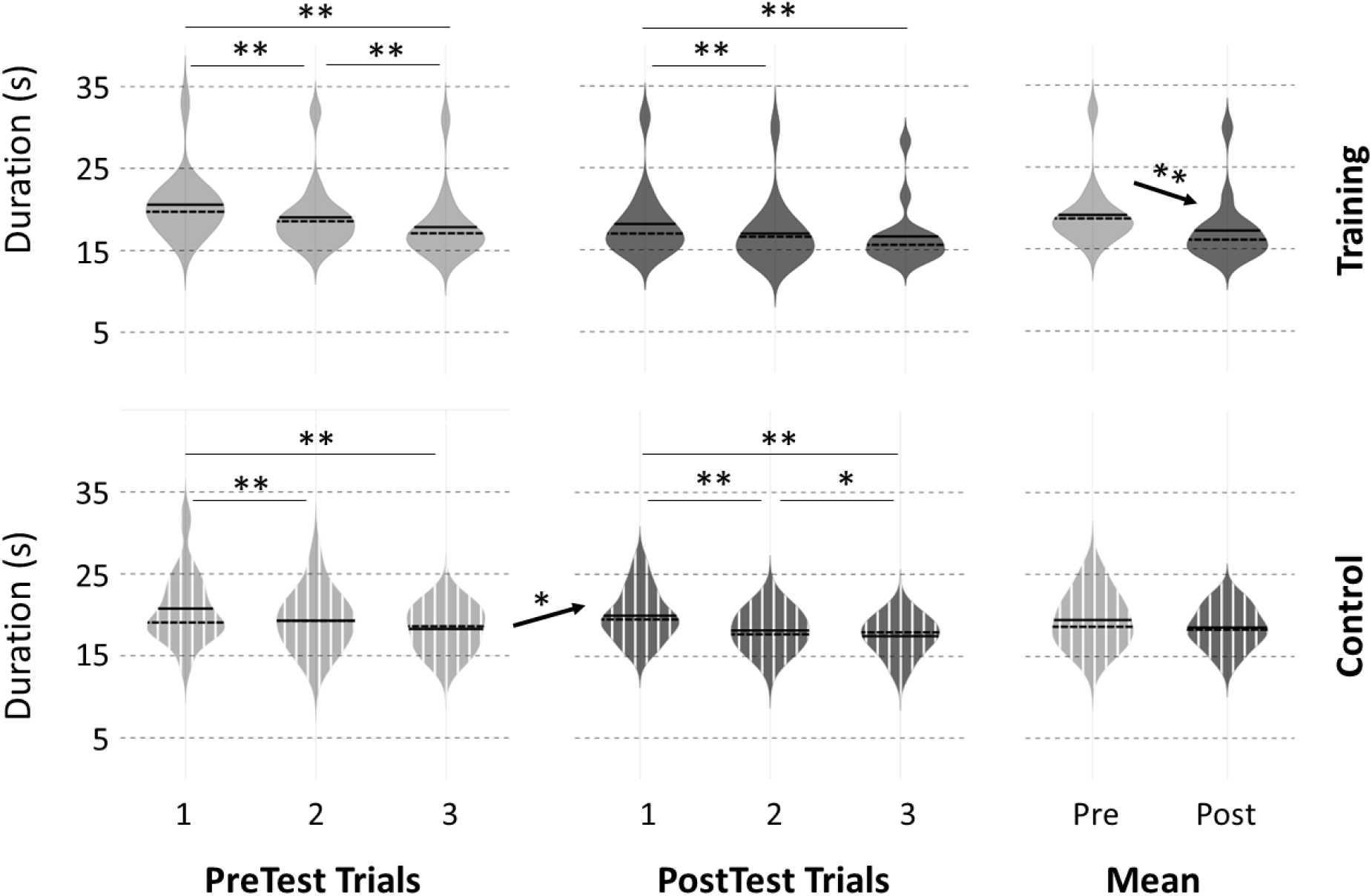
Violin representation of movement duration during the Nine Hole Peg Test for each of the three PreTest and PostTest trials and the mean of the three trials for the mental-training and control groups. Significant difference: *p<0.05, **p<0.01

For both tasks, no significant correlation was observed between the level of improvement and the score for KVIQ, MMSE, TUG or TM6 (all p>0.05).

### Comparison with old adults

The data retrieved from Ruffino et al. (2019) showed that averaged movement duration (NHPT) of old adults decreased from PreTest to PostTest only for the mental-training group (Mental-training, PreTest: 16.6 ± 1.6 s, PostTest: 15.1 ± 2.0 s; Control, PreTest: 19.4 ± 3.2 s, PostTest: 18.5 ± 2.4 s). This effect was marginally significant (Mental-training: p=0.053, t=-2.01), while the Control group did not show any effect (p=0.84, t=0.20). This lack of statistical power after the mental-training could be explained by the small number of participants included in Ruffino et al. (2019). However, the trend is consistent with the one we observed for very old adults.

Old adults showed shorter duration during the PreTest trials compared to very old adults (Mental training, Old: 16.6 ± 1.6 s, Very Old: 19.1 ± 3.9 s, p<0.05, t=-2.18; Control, Old: 13.6 ± 5.7 s, Very Old: 19.4 ± 3.2 s, p<0.01, t=-3.38; Figure 3). After the break in the control groups (PostTest), very old adults were still slower than old adults (Old: 14.1 ± 5.9 s, Very Old: 18.5 ± 2.4 s, p<0.01, t=-2.64). After mental training, the movement duration was not significantly different between old and very old adults (Old: 15.1 ± 2.0 s, Very Old: 17.2 ± 4.0 s, p=0.09, t=-1.73). Furthermore, performance of very old adults after mental training was similar to that of old adults after watching the non-emotional documentary (p=0.12, t=-1.66).

**Figure 3.**
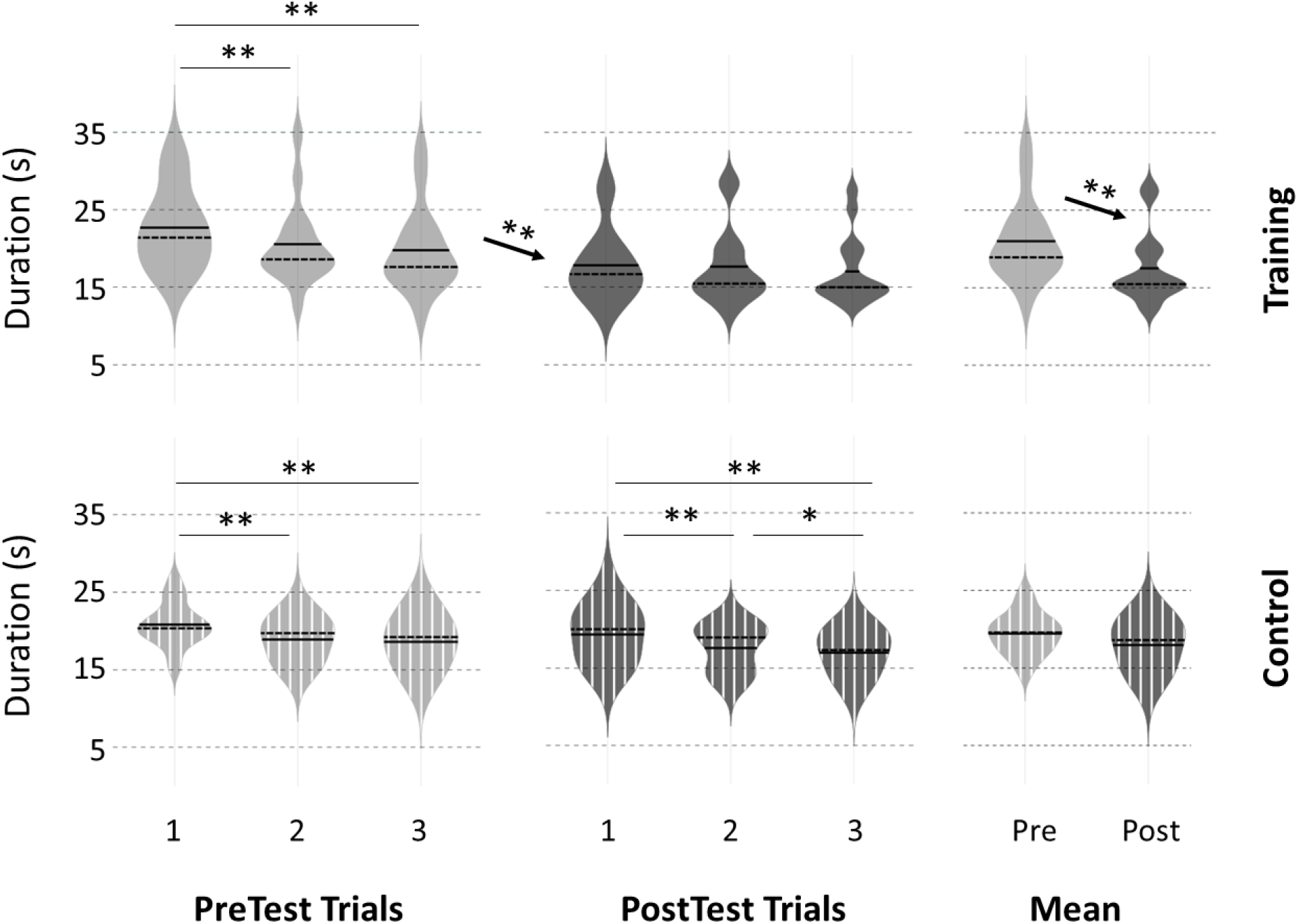
Violin representation of movement duration during the footstep task for each of the three Pre Test and PostTest trials and the mean of the three trials for the mental-training and control groups. Significant difference: *p<0.05, **p<0.01

**Figure 4.**
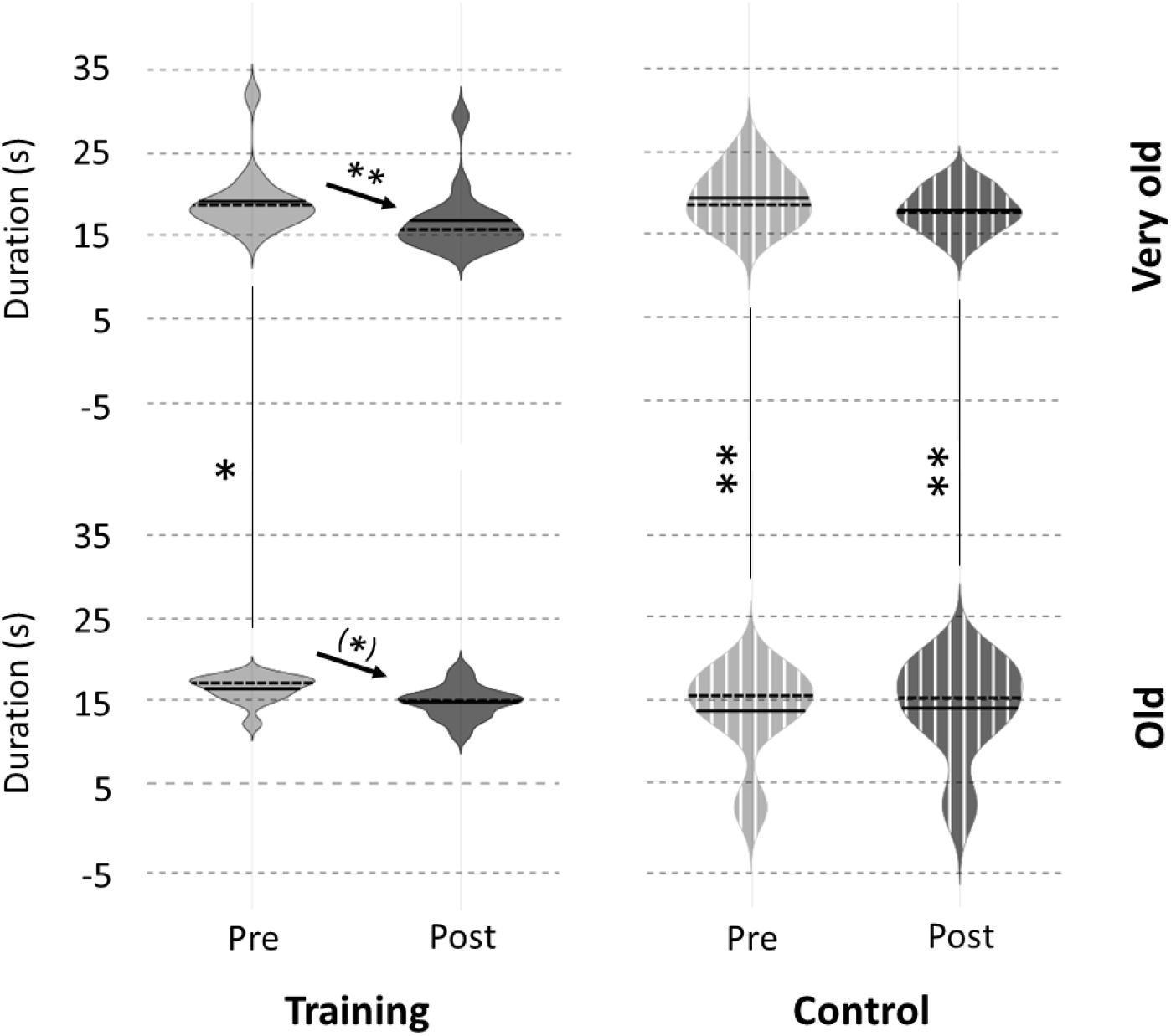
Comparison of movement duration on the NHPT task between our group of very old adults (age: 86 ± 2 years) and a group of old adults (age: 73 ± 5 years) extracted from Ruffino et al. (2019). Means of the three PreTest and PostTest trials are presented for each group. Significant difference: ^(*)^p 0.05, *p<0.05, **p<0.01.

## DISCUSSION

The aim of the present study was to evaluate the effectiveness of motor imagery training on motor memory in very old adults. Although performance was improved after 3 actual trials for both dexterity manual task (upper limbs) and sequential footstep task (lower limbs), the 20min break period (Control group) induced a decrease in performance for the dexterity manual task but no change for the sequential footstep task. However, very old adults who performed a motor imagery training were able to keep their performance improvement for the dexterity manual task, and even increase performance for the sequential footstep task.

### Performance improvement after physical practice

Initial performance at PreTest 1 was similar between the two groups of very old adults, ensuring a homogenous repartition of the participants between the control group and the mental-training group. Thus, we analyzed the improvement of motor performance throughout PreTest trials. The results showed that both mental-training and control groups improved their performance across the three physical trials for NHPT and footstep tasks, with a similar rate of improvement between groups. These results demonstrated that the ability to quickly improve motor performance is maintained at very old age, for both types of task. This finding confirmed previous observations showing that the ability to acquire novel motor skills was maintained with age (Cirillo et al., 2011; Coats et al., 2014). In particular, it generalizes this observation to very old adults and full body balance related movements (footstep task). This is of interest, since negative effects of aging on the neural structure involved in error-based-motor learning have been previously described (e.g. cerebellum; Bickford, 1993). These neurophysiological declines may then affect the ability of seniors to improve their motor performance, only in very particular tasks modalities. It has to be noted that the participants included in the present study were healthy and their functional performance were better than the normative data for community-dwelling people aged over 80 years. Indeed, they walked farther during the 6-min walk test (men: ~455 m, women: ~420m) than the normative data (men: 417 m, women: 392 m (Steffen et al., 2002)) and the duration for the TUG test was similar or lower (men: ~8 s vs. 10 s; women: ~10 s vs. 11s (Steffen et al., 2002)). The sample of participants may not represent the whole population of octogenarians but only the healthiest and the most physically active ones.

### Effects of the 20-min break period on performance

Despite performance improvement after 3 actual trials of NHPT, very old adults returned to their initial performance after a 20-min break, erasing the improvement gains. This alteration in performance after a short break was already observed in old adults for the NHPT (Ruffino et al., 2019), thumb movements (Flöel et al., 2005) or a locomotion task (Malone and Bastian, 2016), whereas young adults were able to maintain their improvement gains. The deficit of motor memory in older adults was proposed to be due to alterations at the cerebral level. More precisely, the primary motor cortex (M1) is mainly involved in the memorization process (Muellbacher et al., 2002; Galea et al., 2011), and some authors showed a dysfunction of this cortical area with aging (Jouvenceau et al., 1998; Sawaki et al., 2003). For exemple, Jouvenceau et al. (1998) observed in aged rat a decrease of NDMA receptors, which play a crucial role for motor memory and synaptic plasticity. Thus, this alteration at cerebral level may partially explain the deficit of motor memorization in older adults. Interestingly, in the present study very old adults were able to maintain their performance at the sequential footstep task after the 20-min break. The better motor memory on the footstep task compared to the NHPT may not only be due to the limbs involved in the task (lower vs. upper limbs) because deficits in motor memory were already observed for the learning of a new walking pattern (Malone and Bastian, 2016). This would rather be due to the difficulty of the task, although we were not able to quantify or rate the difficulty of the two tasks. The NHPT requires fine motor skills to grab the sticks and place them in small holes repetitively, inducing a speed-accuracy tradeoff, whereas the footstep task involved moving the feet in large squares of 0.5-m side, which requires dealing with balance but not precision. Further studies are needed to determine whether the limbs involved in the task or the characteristics of the task influence the performance memorization.

### Effects of mental training on performance

The main finding of the present study is the beneficial effect of motor imagery training on motor memory for both NHPT and sequential footstep task in very old adults. Indeed, for the NHPT motor imagery training allowed the participants to avoid forgetting the performance improvement after a short break. For the footstep task, motor imagery training was even able to improve performance without actual practice. This result is in accordance with previous studies that showed a beneficial effect of motor imagery training on lower limb function of older adults. Indeed, a recent systematic review (Nicholson et al., 2019) provides evidence that motor imagery training improves balance and mobility in older adults. These improvements in motor performance and motor memory may be directly linked with two components. Firstly, at a functional level, motor imagery practice seems to induce positive changes within motor planning (Gentili et al., 2010; Taube et al., 2014). Indeed, via the activation of the neural network implicated in motor planning, motor imagery training could help to refine the internal representation of the considered action by the generation of sensorimotor predictions, leading to an improvement of motor command (Kilteni et al., 2018; Dahm and Rieger, 2019; Ruffino et al., 2022). Secondly, the modulation observed at both neural (Ruffino et al., 2017) and spinal (Grosprêtre et al., 2016) levels during motor imagery seems to reinforce the sensibility and the conductivity of synapses in corticospinal pathway (Avanzino et al., 2015), notably by increasing the effectiveness between pre- and post-synaptic neurons.

### Efficiency of mental training across ages

One of the strengths of the present study is the investigation of very old adults, for whom no data in motor imagery have been previously assessed. Such a terminological separation between old and very old adults may appear artificial, but greater alteration of neuromuscular function occurs after 75 or 80 years old (Venturelli et al., 2018; Varesco et al., 2022). In the present study, although very old adults show similar MMSE score than old adults from the study of (Ruffino et al. 2019), they performed significantly slower the NHPT task. Interestingly, more than being able to limit motor memory deficits, we showed here that mental training was able to compensate the loss of speed between very old and old adults. It has to be noted that to avoid fatigue, the length of the mental training was decreased for very old adults (3 blocks of 10 trials compared to 5 blocks of 10 trials for old adults (Ruffino et al., 2019)). Nonetheless, this shorter training duration was sufficient to improve performance. Taken together, these results suggest that a brief session of motor imagery (~20 min) could be an efficient and low cost rehabilitation method for very old adults.

## Conclusion

The present study extended the benefits of motor imagery training to the very old population, showing that even a short motor imagery training improve their performance and favor the motor memory process. These results confirmed that mental training, using motor imagery, is an interesting alternative option to complement traditional rehabilitation protocols. Mental training characteristics should be optimize in the future to maximize the beneficial effect of motor imagery in very old adults.

